# Nucleation of Multiple Buckled Structures in Intertwined DNA Double Helices

**DOI:** 10.1101/196345

**Authors:** Sumitabha Brahmachari, Kathryn H. Gunn, Rebecca D. Giuntoli, Alfonso Mondragón, John F. Marko

## Abstract

We study the statistical-mechanical properties of intertwined double-helical DNAs (DNA braids). In magnetic tweezers experiments we find that torsionally-stressed stretched braids supercoil via an abrupt buckling transition, which is associated with nucleation of a braid end loop, and that the buckled braid is characterized by proliferation of multiple domains. Differences between the mechanics of DNA braids and supercoiled single DNAs can be understood as an effect of increased bulkiness in the structure of the former. The experimental results are in accord with the predictions of a previously-described statistical-mechanical model.

---

Catenated or intertwined DNA molecules are a common occurrence in the cell as they are an intermediate in segregation of sister chromatids, following DNA replication and recombination [1–5]. Catenated DNA molecules can be mimicked *in vitro* by wrapping or “braiding” two single DNA molecules around each other. At the single-molecule level, DNA braids are important substrates to study the topology-manipulation mechanism of DNA topoisomerases and site-specific DNA recombinases [5–8].

Despite its importance, our understanding of the mechanics of DNA braids lags behind that of twisted single DNA molecules, where precise single-molecule experiments have successfully characterized the supercoiled state and its nucleation, driven by torsional stress in the duplex DNAs [9–15]. Qualitative and quantitative theoretical predictions have played a key role in our unstanding of the mechanics of single supercoiled DNAs [16–20], but are lacking in the case of braided DNAs. In this Letter, we study braiding of two freely-swive duplex DNAs, and find that the mechanical prope of braids significantly contrast with those of single percoiled DNAs, largely due to the increased *struct bulkiness* (i.e., higher bending stiffness and excluded ume) and the variable twist rigidity of braided DNAs.

Previous experimental studies on nicked-DNA br [21–23] have revealed a change in slope of the braid extension versus catenation curve [also seen in Fig. (1b)]. Monte-Carlo simulations [22, 23] similarly suggested formation of a plectonemically supercoiled braid, however, the characterization of the buckled state was incomplete. Here, we study braided torsionally-relaxed DNA double helices, and we observe an *abrupt* nucleation of a buckled braid state followed by proliferation of multiple domains in the plectoneme-coexistence state, in accord with theoretical predictions [24]. Our work highlights the significance of structural bulkiness in buckling of DNA, which may have relevance in understanding the mechanical response of bulkier DNA structures such as protein-coated DNAs.

**FIG. 1:**
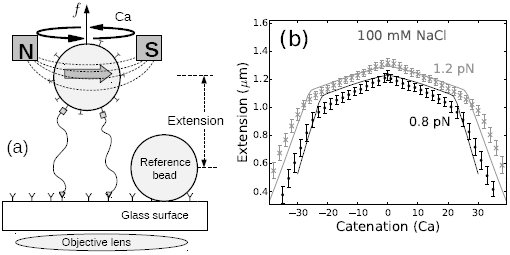
(a) Schematic of the magnetic tweezers setup. Two DNA molecules are attached (using only one of the DNA strands as attachments) to the glass surface and a paramagnetic bead via digoxigenin-antidigoxigenin and biotin-streptavidin interactions, respectively. The single-strand attachments ensure no twisting of individual DNA molecules upon rotation of the bead [8, 25]. (b) Experimentally measured end-to-end extension of a braid as a function of catenation number for 0.8 (black points) and 1.2 pN (gray cross marks) applied forces in 100 mM NaCl, where the error bars represent standard deviation. The solid lines represent theoretical predictions from Ref. [24] for 1.6 *μ*m DNA molecules, with intertether distance 0.19 *μ*m, and under 0.8 (black line) and 1.2 pN (gray line) force at 100 mM monovalent salt. The small peak in extension at zero catenation is due to the small intertether distance, *i.e.*, the close proximity of the two DNA molecules for this particular DNA pair [24].

We used bright-field magnetic tweezers [25] to study braided DNAs, where we attached one pair of ends of the two double-helical DNAs to a glass surface and the other pair of ends to a one-micron paramagnetic bead [8] [Fig. (1a)]. The inter-DNA linking (or “catenation”) number in the braid is controlled by rotating the bead using the magnet, whereas the applied force is controlled by varying the distance between the magnet and the bead. To ensure that each DNA is not subject to doublehelix twisting torque, only the 5^*/*^-ends of the DNAs were attached to the surfaces, allowing swiveling of the free strand about the tethered one, which makes the number of turns of the bead a direct measure of catenation in the braid. All experiments were carried out in 100 mM NaCl buffer.

As the magnet is rotated, the extension of the braid un der fixed force decreases when the catenation is increased, producing a characteristic bell-shaped curve [Fig. (1b)]. Fig. (1b) shows a typical plot of braid extension as a function of catenation for 0.8 and 1.2 pN forces under physiological salt conditions (100 mM NaCl, 20 mM TrisHCl, pH 8); solid lines are theoretical predictions of the model in Ref.[24]

The change in extension of the braid between catenation numbers 0 and 1 is related to the distance between the two tether points; closely-spaced DNA tethers produce a small jump, whereas, a larger separation between the DNA tethering points show a sharper initial jump [21–24]. In Fig. (1b), the jump is small relative to the length of the braided molecules (≈1.6 *μ*m), indicating that the two DNAs are tethered close together (*≈* 0.19 *μ*m). After the first catenane is introduced, the extension of the braid decreases with increasing catenation due to formation and consequent increase in size of the helically-wrapped region of the braid. Since the individual DNA molecules cannot be supercoiled, the extension plots are symmetric for positive and negative catenations [Fig. (1b)]; our model assumes this symmetry as there are no DNA-twist-energy terms in the free energy expressions[24].

The predicted torque in the braid increases non-linearly with increased catenations, implying a catenation-dependent twist modulus in braids [23, 24, 26–28]. This marks a significant difference between the mechanics of braided DNAs and twisted single double-helix DNAs, where a constant twist stiffness results in a remarkably linear torque response [10–12].

The downward slope of the extension plot steepens at the point where a braid plectoneme nucleates, and the extension continues to decrease sharply past the nucleation of the buckled phase [Fig. (1b)]. Buckling releases torsional stress in the braid due to the inter-braid writhe contribution to the total catenation number. The braid plectoneme also contains a “teardrop”-shaped braid loop (end loop), where the braid bends by 180 degrees [Fig. (2b)]. The energy associated with formation of the end loop acts as a nucleation energy cost of a plectoneme domain.

**FIG. 2:**
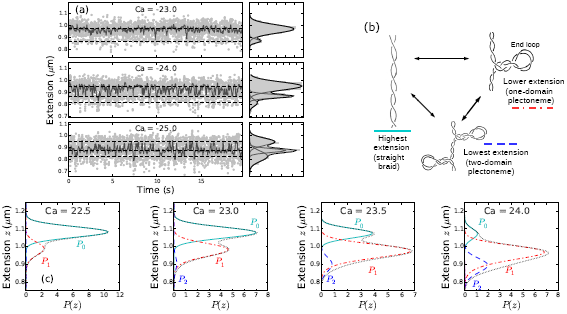
(color online) (a) Time series of end-to-end extension of a braid constituted of two torsionally-unconstrained 6 kbDNAs, under 0.8 pN force at100 mM NaCl [Fig. (1b)], for three catenation numbers (Ca = −23, −24, and −25) near the buckling transition point [the point of slope change in the extension curve, Fig. (1b)]. Data were collected at 200 Hz (light gray dots), then median filtered (dark points) using a 0.1 sec time window to show dynamic switching between discrete extension states. The panels on the right of each time series plot show histograms of the raw data using 10 nm bins (gray shaded area); the y-axis is the same for left and right panels. The histograms were fit to a sum of multiple Gaussian distributions, where the dark line is the best-fit distribution and corresponds to the sum of the individual Gaussians shown in gray lines. The sum of two Gaussian distributions fit the data better at low catenation (Ca=-23), indicating nucleation of the first plectoneme domain, whereas, the sum of at least three Gaussians is required to fit the histograms at higher catenation (Ca= −24, −25), due to the appearance of multiple plectoneme domains. (b) Schematic diagram of three braid-extension states accessible via thermal fluctuations near the buckling transition. (c) Theoretically predicted (using the model in Ref. [24]) extension histograms near the buckling transition, where the black dotted line shows the total extension distribution (*Ptot*); also plotted are the individual contributions from the straight braid (*P*0, cyan solid line) and the buckled braid with one (*P*1, red dot-dashed line) and two (*P*2, blue dashed line) plectoneme domains [see Eq. (3)]. Contributions from the three (*P*3) and the four-domain (*P*4) plectonemes are also plotted, however, the negligible statistical weight of those states for the plotted range of catenation renders them almost invisible in the predicted histograms.

Appearance of a plectoneme domain requires nucleation of the braid end loop, which causes a discrete change in braid extension since the end loops are finite-sized structures. Fig. (2a) shows data for a time series of braid extension under 0.8 pN force and 100 mM NaCl salt concentration, at a fixed catenation near the buckling transition, *i.e.*, near the point of slope change in the extension curves [Fig. (1b)]; and the histograms show the probability density of braid extension. Near the buckling transition point [Ca=-23 at 0.8 pN, see Fig. (1b)], the probability distribution of braid extension is bimodal [Fig. (2a)], where the higher and the lower extension peaks respectively correspond to the straight braid and the one-domain plectoneme braid (*i.e.*, a plectoneme with one end loop) [Fig. (2b)].

In the vicinity of the buckling transition, the occupancy of the lower-extension state increases with increasing catenation, due to the appearance of the first buckled plectoneme domain. Simultaneously, the occupancy of the higher-extension state decreases as the purely-straight state of the braid disappears. The data also show appearance of multiple discrete-extension states after the nucleation of the first domain [multiple peaks in the histograms for Ca= −24 and −25, see Fig. (2a)], where the lowest-extension state corresponds to a two-domain plectoneme braid [plectoneme with two braid end loops, see Fig. (2b)]. Braids being bulky structures favor multiple small plectoneme domains over a single large one, where structural bulkiness is derived from bending stiffness and excluded diameter of the braids. Since braids have two wrapped double helices, the effective braid bending stiffness is twice that of a double helix. Also, due to the electrostatic interactions, braids have an excluded diameter which is at least twice of that of a single double helix. Larger excluded volume increases the lower bound on braid-plectoneme diameter, which destabilizes the superhelical state relative to the braid end loops [24]. Increased bulkiness makes the two-domain plectoneme structure fluctuation-accessible, and results in the appearance of a finite-probability state with extension lower than that of the one-domain plectoneme. The probability of occupancy of the one-domain plectoneme state increases past the onset of the buckling transition, and then decreases as the two-domain state becomes more probable. From the median-filtered time signal [Fig. (2a)], we estimate the nucleation rate of a braid plectoneme state ≈10 s^−1^ which is similar to that observed for a plectoneme domain in twisted single dsDNA [14].

Following the theoretical work in Ref. [24], we consider DNA braids featuring a coexistence of straight and plectonemically buckled states, where every plectoneme domain is accompanied with a loop-shaped braid [Fig. (2b)]. The braid end loop acts as a nucleation cost to a plectoneme domain, which we estimate as the total elastic energy associated with forming the end loop: *βE* = 2*εA/*γ + *βf* γ. The first term is the total bending energy of a teardrop-shaped loop (*ε* = 16) [29, 30], with the bending persistence length of DNA *A* = 50 nm, and the size of each braiding strand in the loop γ. The second term is the work done in decoupling the braid loop from the external force *f*; we define *β* ≡ 1*/k*_B_*T* (*T* = 290*K*).

Minimizing the total energy of the loop with respect to γ, we find the equilibrium length of the end loop associated with each plectoneme domain:

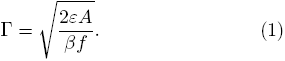

The presence of a finite-sized (γ) end loop causes an abrupt change in braid extension upon nucleation of a plectoneme domain. We construct a canonical partition function *Ƶ* to include equilibrium fluctuations among states with various plectoneme lengths *L*_*p*_, and number of domains *m* [24]. The extension distribution in each of the summed-over state is Gaussian:

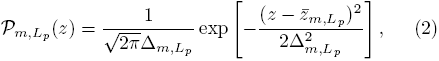

where the mean and the variance are respectively given by: 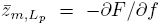 and 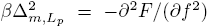. Here, *F* (*L*_*p*_*, m*) is the total free energy of the plectoneme-coexistence state, which includes the force-coupled straight braid, superhelically-bent braid of size *L*_*p*_, and *m* end loop(s) [24]. The total distribution of extensions at a given catenation is obtained from summing all the corresponding Gaussian distributions [Eq. (2)] with their respective Boltzmann weights:

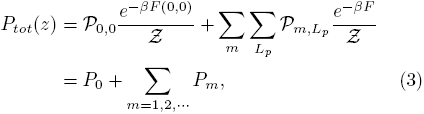

where *P*_0_(*z*) and *P*_*m*_(*z*) are the respective contributions to the total extension probability distribution *P*_*tot*_(*z*) from the straight and the *m*-domain plectoneme states.

Fig. (2c) shows the predicted probability distributions of extension near the buckling transition point for braids made up of 1.6 *μ*m DNAs separated by 0.19 *μ*m (*d* = 0.12*L*) at the tethering points, in 100 mM monovalent salt condition and under 0.8 pN force [Fig. (1b)].

The total probability distributions *P*_*tot*_ [black dotted lines, Fig. (2c)] are multimodal, similar to the experimentally observed histograms. The individual Gaussian contributions from the straight phase (*P*_0_), and the plectoneme phase with one (*P*_1_) and two (*P*_2_) domains are also plotted in Fig. (2c). Near the buckling point, increasing the catenation in the braid makes the purely-straight braid (*P*_0_) less favorable than the one-domain plectoneme-coexistence state (*P*_1_). Further increase in catenation number leads to the nucleation of new plectoneme domains, which gives a strong asymmetric character to the extension distributions. In Fig. (2c), the contributions corresponding to three (*P*_3_) and four-domain (*P*_4_) plectonemes are also plotted, however, they are almost invisible due to negligible statistical weights of the states.

Fig. (3) shows the comparison of theoretically predicted change in extension upon nucleation of the first and the second domain of braid plectoneme with experimental data. For both theoretical calculations and experimental measurements, we define the extension jump upon nucleation of the first plectoneme domain as the distance between the means of the extension distributions corresponding to the straight (*P*_0_) and one-domain braid plectoneme (*P*_1_), when both the states are almost equally-likely and most-probable. Similarly, the extension jump associated with nucleation of the second plectoneme domain is defined as the distance between the means of the extension distributions corresponding to one (*P*_1_) and two-domain plectonemes (*P*_2_), one catenation unit after the first jump is measured.

**FIG. 3:**
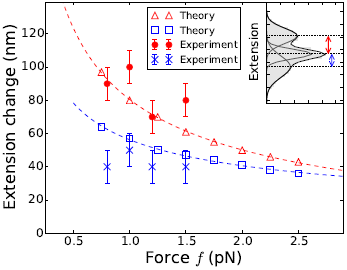
(color online) Comparison of theoretically-predicted change in extension upon nucleation of the first (red open triangles) and the second (blue open squares) plectoneme domain with experimental observations. Red points and blue cross marks represent the difference in extension between successive peaks (inset) in the experimental best-fit histograms, where the second jump (blue cross markers) is measured one unit catenation after the first jump (red points). The error bars represent binning error. The red and blue dashed lines show the expected *f*^-1/2^ power law [Eq. (1)], best-fit to red open triangles and blue open squares, respectively.

The predicted magnitude of extension jump upon nucleation of a plectoneme domain decreases with increasing force due to the decrease in size of the nucleated braid end loop [Eq. (1)], although this trend is not apparent in the experiments [Fig. (3)]. However, we find both theoretically and experimentally that the extension change associated with nucleation of the second plectoneme domain is significantly smaller than that of the first one, suggesting that the nucleation energy cost of the first domain is larger than that of the second domain.

In conclusion, the results show that the response of stretched DNA braids under torsional stress is distinct from that of twisted single DNA molecules. In both the cases, applied torque drives a buckled plectoneme structure [12, 13, 21–23], nucleation of which is associated with a discontinuity in the extension. However, in the case of supercoiled single DNA molecules in physiological salt conditions (≈100 mM Na^+^), injection of twist in the buckled state mainly leads to the growth of a single plectoneme domain, whereas, in the case of braids, many short plectonemes appear after the buckling transition. The relatively large curvature energy associated with plectonemic buckling of braids makes the superhelical structure less favored compared to that in single supercoiled DNAs. This is roughly analogous to behavior of supercoiled single DNAs at low salt conditions (≈10 mM Na^+^), where DNA excluded volume is enhanced and where multiple small plectonemes are found [14, 15, 18, 20].

Our results also suggest that stretched bundles of DNA under torsional stress are more likely to form multidomain “looped” structures than long plectonemes due to the increased bulkiness; however, close packing of the DNAs in a bundle may lead to novel structural properties. Similar effects of bulkiness may also appear in mechanics of protein-covered DNAs. Overall, the contrast between the mechanical properties of braided and supercoiled DNAs can be simply interpreted as a result of the structural bulkiness and linking-number-dependent elastic moduli in braids, and is well explained by the theoretical model [24]; whether these potential differences directly influence cellular function or the way proteins interact and modify DNA remains to be determined.

The authors acknowledge support from the National Institutes of Health (Grants R01-GM105847, R01-GM05135, U54-CA193419 [CR-PS-OC], NRSA pre-doctoral training grant T32-GM008382, and subcontract to Grant U54-DK107980), and by the National Science Foundation (Grants DMR-1206868 and MCB-1022117).K.H.G. also acknowleges support by a Dr. John H. Nicholson Fellowship.

